# Opposing responses of hippocampal theta oscillations to running and a forelimb-dominated sensorimotor behavior

**DOI:** 10.1101/2025.06.08.658384

**Authors:** Gabe Holguin, Krystina Jorgensen, Andrew Tapia, Gianna A. Jordan, Abhilasha Vishwanath, Carol A. Barnes, Stephen L. Cowen

## Abstract

Hippocampal theta oscillations regulate the timing of neurons to support navigation, memory formation, and sensorimotor integration. Theta is modulated by running speed, breathing, whisking, and jumping and increases in tasks involving memory encoding or retrieval. The positive relationship between theta frequency and running speed is believed stabilize hippocampal representations of space amid movement variability. Here, we incorporated a novel string-pulling task to determine if established relationships between movement and theta hold when progress to a reward is determined by the length of string pulled. This task eliminates many speed-associated inputs, such vestibular, visual, and hindlimb information, and allows an unprecedented level of precision in the analysis of individual paw movements. Given that animals move the string a fixed length to acquire a reward, we predicted that the positive relationship between theta frequency and speed would hold.

**Approach:** Five Sprague Dawley rats (4 mo.) were trained to continuously pull a string a fixed distance of 208 cm using an automated string-pulling system and run on a track for food reward. Local-field data was acquired from electrodes in dorsal CA1.

**Results:** Relationships between theta and movement speed were distinct during string pulling and running. While theta was robust in both conditions, frequency was significantly reduced during string-pulling and showed no speed-frequency coupling, unlike running. This difference could result from the conflict between hindlimb and forelimb signals, with only forelimb movement signaling advancement. Fine-grained analysis of paw movements during string-pulling (lift, advance, grasp, pull, push) revealed that theta power and frequency peaked during the contralateral paw’s downward push despite paw speed being low during this action. This suggests that theta frequency and power could respond to effort rather than purely kinematic information. Notably, running-associated theta may similarly reflect both speed and effort as most locomotor tasks conflate these variables. Finally, theta phase aligned from one reach-pull cycle to the next during the downward pull motion - the first action that directly advances the string forward. Since phase-locking has been associated with sensorimotor gating, synchrony at this point could reflect the gating of inputs that are the most causally relevant for reaching the reward, potentially facilitating integration of action-outcome signals for memory encoding and navigation. Taken together, these data support a dual-scale view of hippocampal processing and theta-band activity where macroscale theta activity requires suprathreshold sensory, vestibular, and proprioceptive drive and microscale theta remains sensitive to subsecond limb movements.

## Introduction

Hippocampal theta oscillations are a signature of active exploration and voluntary movement in rodents, with their frequency and power closely linked to running speed and spatial navigation^1,2^. This relationship is thought to facilitate the stabilization of spatial representations and support episodic memory formation by temporally organizing neuronal firing patterns^3–5^. While the modulation of theta by gross locomotion speed has been extensively characterized, comparatively less is known about how theta dynamics are influenced by fine motor actions, particularly those involving skilled forelimb movements in goal-directed tasks.

To address this gap, we investigated hippocampal theta activity during a string-pulling task in rats, a behavior requiring precise, rhythmic bimanual forelimb coordination and proprioceptive guidance in the absence of direct visual feedback. String-pulling has been used to assess behavior in more than 160 species^6^, including humans^7^, and has been recently used for the investigation of motor control, movement disorders, and stroke^8–10^. A common version of the task requires animals to pull a fixed length of string to receive reward. Thus, this task is goal oriented and draws parallels to spatial navigation tasks where distance traversed, instead of string length pulled, predicts reward delivery. By comparing neural activity during string pulling and track running, we sought to determine whether the established positive relationship between theta frequency and movement speed extends to skilled, non-locomotor behaviors. Furthermore, this task, as implemented here^11^, eliminates many speed-associated inputs, such vestibular, visual, and hindlimb information. Given that animals move the string a fixed length to acquire a reward, we predicted that the positive relationship between theta frequency and speed would hold.

Another feature of the task is that it allows detailed measurement of individual paw movements and thus allows the investigation of microscale actions as well as macroscale measurement of movement speed. This is important as some of the earliest investigations of theta activity noted increased frequency and power during object manipulation^12^ and to the magnitude of an anticipated jump^12,13^. Theta phase has been shown to align to individual actions such as button-presses^14^, eye saccades^15^, whisking and breathing^16,17^, and footsteps^18^, and often when such actions are during tasks involving memory encoding or retrieval. Consequently, we predicted that theta power, phase, and frequency would synchronize to isolated motor components of the reach-grasp-pull movement.

Our results reveal a striking dissociation in theta dynamics between running and string-pulling. Although robust theta oscillations were present in both behaviors, theta frequency and its harmonic were significantly reduced during string-pulling compared to running, and the canonical positive correlation between movement speed and theta frequency was abolished during string-pulling. Fine-grained analysis demonstrated that theta frequency and power peaked during the ‘push’ phase, when paw speed was lowest. In addition, theta phase-locking was maximal during the ‘pull’ phase, the phase that first advanced the string towards the reward.

These findings suggest that hippocampal theta integrates proprioceptive and effort-related signals in a task-dependent manner, and that its temporal structure is finely tuned to the sensorimotor demands of skilled, goal-directed actions.

## Materials and Methods

### Animals

Five male Sprague Dawley rats (∼3-4 months old, weighing 360-420 g, Envigo RMC Inc., Indianapolis, IN) were single-housed in a temperature and humidity controlled 12-hr reverse light/dark cycle room with food and water available *ad libitum* during the habituation period. During behavioral training, rats were food restricted to 85% of their *ad libitum* body weight. All procedures were in accordance with NIH guidelines for the Care and Use of Laboratory Animals and approved IACUC protocols at the University of Arizona.

### Track-running and string-pulling behavioral training

Prior to electrode implantation, animals were trained to run on a circular track and pull strings. <u>Track-running training:</u> Rats were trained for ∼3 days to run clockwise on a 54 cm diameter circular track and animals received Ensure® food reward after each lap. Criterion performance was reached when rats ran at least 20 laps per training session. <u>String-pulling</u> <u>training:</u> Training stages are summarized in **Figure 1G**. Stage 1 of training involved tying a Cheerio® to the end of a cotton string and draping the string over the wall of the enclosure and animals were left in the box for ∼30 minutes to explore and pull the string. Strings were initially ∼12 cm and the length was progressively increased in ∼30 cm increments until rats consistently pulled ∼100 cm. Once they reached criterion, animals were shifted to the automated string-pulling apparatus (Stage 3) where rewards were automatically dispensed when the designated length was pulled. Criterion performance was met when animals consistently pulled 208 cm for food reward.

**Figure 1.**
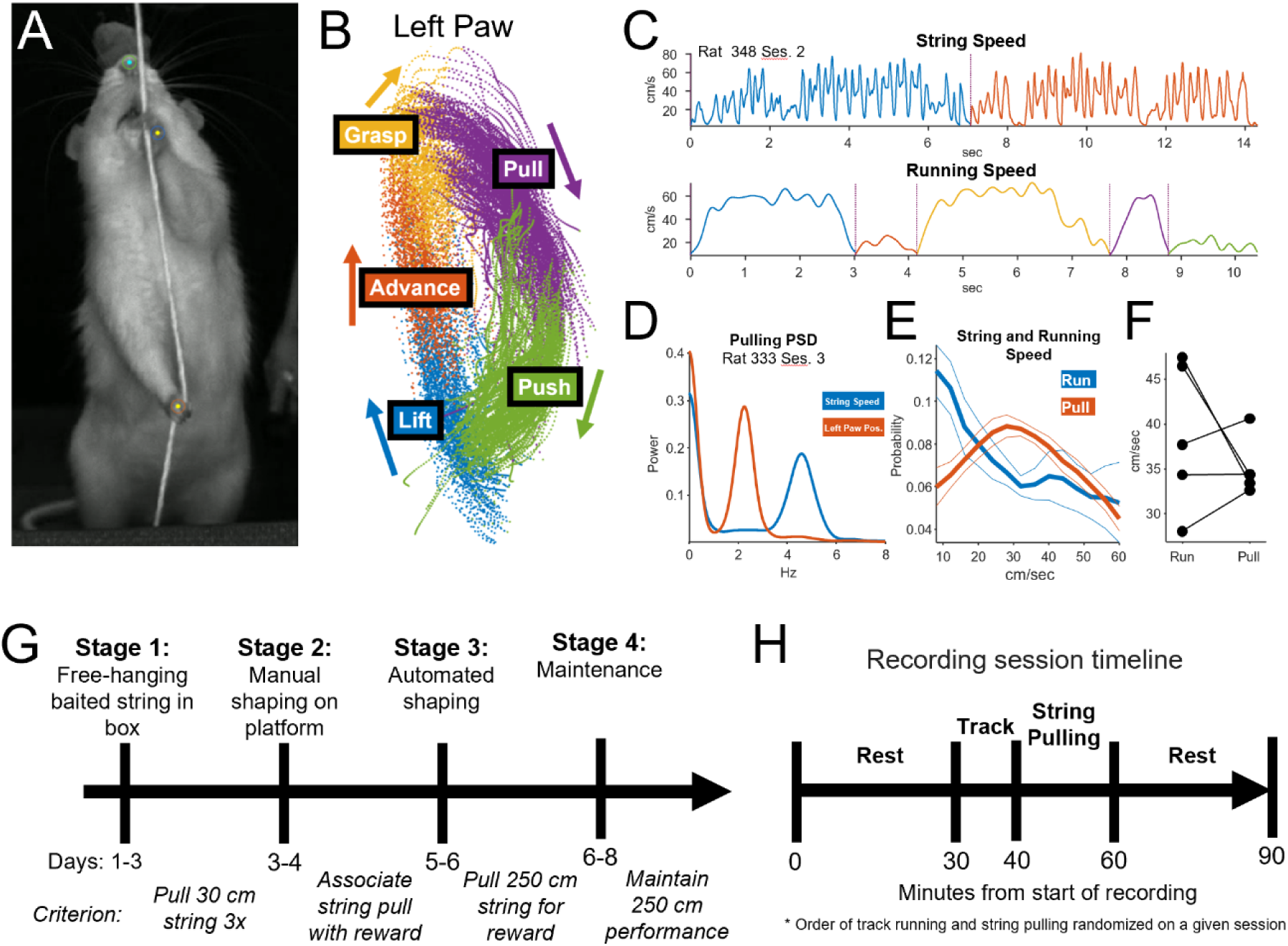
Animal Behavior. A) Still photo from video acquired during string-pulling. Dots indicate tracking of the nose and paws (DeepLabCut). B) Trajectory of the left paw throughout a single behavioral session. Colors indicate the results of automatic segmentation of the 5 phases of string pulling (lift, advance, grasp, push, pull). C) Top: Plot of string speed during two pulling bouts (colors). The vertical line indicates the separation of the bouts (the bouts were not contiguous). Bottom: Running speed during 5 trials (colors) on the circular running track from the same session. D) Power-spectral density of the time-course of string speed (blue) and paw movement (orange) during a single session. The peaks indicate that the string speed fluctuated at ∼5 Hz while the left paw was half of this (2.4 Hz). E) Mean ± SEM distribution of speeds for track-running (blue) and string-pulling (orange). F) While the shape of the distributions of speed differed during track running and pulling (E), a paired t-test indicated no difference in the mean speed per rat (p = 0.39, n = 5). G) The timeline for training the string-pulling behavior. H) The timeline for a given neural recording session. The order of track running and string pulling was randomized on each session.

### String-pulling apparatus

The Pulling And Neural Data Analysis (PANDA) system^11^ consists of a loop of string mounted on pulleys with one pully connected to a digital rotary encoder (BQLZR, Shenzhen, China). The encoder provides precise measurement of string speed, and the signal was monitored by a connected Arduino™ microcontroller. The microcontroller also controls the delivery reward when a designated length of string is pulled or if the user manually triggers delivery. Reward is delivered through a solenoid valve and spout located 30 cm from the string. Once animals reliably pulled the string on the PANDA apparatus, rewards were delivered automatically at predetermined lengths of string pulled. Details regarding the construction and animal training can be found in Jordan et al.^11^, and all code is available on GitHub (https://github.com/CowenLab/String_Pulling_System).

### Surgical procedures

Rats were anesthetized using isoflurane (1 - 3% isoflurane mixed with oxygen, 1.5 L of oxygen/min). Isoflurane was lowered after induction and implantation until the animal’s breathing stabilized at ∼50 breaths per minute. Rats were implanted with custom 32-channel tetrode arrays. Each array consisted of 8 twisted-wire tetrodes constructed of 12 µm diameter polyamide-insulated nickel chromium steel wire. A shorted reference and ground wire soldered to a skull screw was placed over the cerebellum. The tetrode array was implanted in CA1 in the right hemisphere (AP: -3.0, ML: +2.0, DV: -2.5). Analgesia and antibiotic ointment over the surgical incision were administered for two days following surgery. Animals recovered for at least one week prior to neural recording. Procedures were performed in accordance with National Institutes of Health guidelines for laboratory animals under protocols approved by the University of Arizona Institutional Animal Care and Use Committee.

### Electrophysiology and experimental procedures

Electrophysiological data was acquired at 30 kHz using the Intan neural recording system (Intan Technologies Inc.). Local-field recordings were subsequently downsampled to 500 Hz for analysis. The timeline for an individual neural recording experiment/session is presented in **Figure 1G**. Each session began with 30 minutes of baseline measurement as animals rested in a pot. Rats then performed track-running and string-pulling behaviors with the order of these behaviors being counterbalanced across recordings sessions. During track-running, the animal ran clockwise on a 54 cm diameter track for 10 minutes. During string-pulling, rats performed the string-pulling task described previously on an elevated platform (**Figure 1**) for 20 minutes. After completion of these behaviors, the rat was again allowed to rest for 30 minutes in a towel-lined pot. Each rat completed ∼5 sessions over a two-week period.

### Motion tracking

A side-facing camera (MakoU130b, Allied Vision, Stadtroda, Germany) monitored string-pulling behavior and data was acquired at 367 fps using the Image Acquisition Toolbox in Matlab (2019). A top-down camera (Manta G-033C, Allied Vision, Stadtroda, Germany) monitored track running and data was acquired at 60 fps using NorPix software (Montreal, Quebec). DeepLabCut^19^, was used to track the nose and paw position during string pulling and the nose, ear, and tail position during track running.

### Behavioral Segmentation

Each pull of the string was automatically segmented into reach and withdrawal phases as well as the fine-grained “lift, advance, grasp, pull, and push” phases (**Figure 1B**). These phases were comparable to those described in Blackwell *et al.,*^9^. Segmentation was accomplished through a three-step process. First, the time series of vertical (y) paw data (sampled at 367 Hz) was band-pass filtered (1-8 Hz). The Hilbert transform of this output yielded a continuous measure of pull phase (Matlab: angle(hilbert(y data))). Second, the velocity (positive up, negative down) and acceleration were determined from the x and y time-series data. Third, data from each paw was analyzed cycle-by-cycle to identify the categorical label of each pull phase (lift, advance, grasp, pull, push). For example, the transition from “pull” to “push” was determined as the time when the paw position during a downward motion (negative velocity) reached its maximum eccentricity in the x dimension. Segmentation accuracy of the automated procedure was confirmed visually by comparing videos of each reach or withdrawal to the output of the automated procedure.

### Histology

Upon completion of ∼5 recording sessions, electrolytic lesions were made at each recording site to confirm targeting. Three days following the lesion, animals were sacrificed using 0.35 ml euthanasia solution (390 mg/ml pentobarbital sodium and 50 mg/ml phenytoin sodium; Vetone, Boise, ID) and perfused with 4% paraformaldehyde in phosphate-buffered saline. Brains were extracted, stored in 30% sucrose solution, and coronally sectioned. Tissue was stained using cresyl violet for verification of electrode placement. Electrolytic lesions were found in all but one rat – this animal was included in the analysis given clear evidence for tetrode track damage above the CA1 region and robust theta activity during running and pulling behaviors (**Figure S1**).

### Data analysis

#### Local field potential (LFP) Analysis

LFP signals from CA1 were referenced to the cerebellar skull screw signal, and the acquired signal was downsampled (using resample in Matlab) from 30 kHz to 500 Hz for analysis. Theta (6-12 Hz) and its harmonic (12-24 Hz) were isolated using a 12^th^ order Butterworth band-pass filter (designfilt). Instantaneous power and phase within the target band were determined using the Hilbert transform. LFP was only analyzed for epochs where animals were moving continuously between 10 - 70 cm/s. For the analysis of acceleration/deceleration, epochs were limited to ±25 cm/s². Wavelet transforms of the LFP data were performed using a bump wavelet (cwt()) as the bump wavelet has a higher frequency resolution than the Morelet wavelet.

#### Statistical Tests

Inferential tests used the animal as the sample (n = 5 rats). To acquire a measure from a given animal, values of the variable in question (e.g., the correlation between theta and running speed) were averaged across sessions for a given animal (∼3 sessions per animal). This value was then used as the within-animal measure. Unless otherwise stated, paired- and one-sample t-tests (α = 0.05) were used to assess statistical significance. Statistical analyses were performed using R-Studio 2023.

## Results

### Behavioral performance

#### Pre-training

Prior to microdrive implantation, each rat was trained to pull a minimum of 2 meters of string per pulling bout and was required to complete >10 bouts per 30-minute session. A bout was defined as a period where the rotary encoder indicated motion of the string for ≥1 second. A bout ended when the rotary encoder was immobile for ≥1 second. Rats required <8 days to meet the 2-meters-per-bout criterion. **Figure 1G** summarizes the pre-training procedure.

#### Behavior during recording sessions

Each session consisted of a block of string pulling and a block of running on the circular track (**Figure 1H**). During neural recording, animals achieved at least 42 string-pulling bouts per session, with individual bouts lasting ∼5.5 seconds with a total length of string pulled per session approaching 90 meters (**Figure S2**). Examples of string speed and running speed for a given session are shown in **Figure 1C** with the color indicating individual pulling or running bouts. The power spectral density in **Figure 1D** shows the strong ∼5-Hz rhythmicity in string speed and ∼2.5-Hz rhythmicity in left paw speed from data acquire and frequency would synchronize during a single session. Paw trajectories were categorized into lift, advance, grasp, pull, and push phases (see **Figure 1B**, see Methods). While the shapes of the distributions of string speed and running speed differed (**Figure 1E**), there was no statistical difference between the mean pulling and track running speed (**Figure 1F**; paired t-test, p = 0.39, n = 5 rats).

### Theta oscillations were robust during string pulling but at a lower frequency and with a notably reduced harmonic relative to track running

While theta has been reported during bimanual behaviors, such as the manipulation of food pellets^12^, it had not yet been reported during string pulling. String-pulling differs from bimanual object manipulation in a few ways. First, during string pulling, rats look away from the string and thus do not use vision to manipulate the string, presumably relying on proprioceptive cues to guide the string to the contralateral paw within a given pull cycle. Second, the string-pulling task described here is goal-directed in the sense that the length of string pulled determines when the reward will be delivered which was somewhat homologous to running on a track where the distance traversed typically predicts reward delivery.

Analysis of the spectral response of the LFP (Welch’s power spectral density (PSD)) during track-running and string-pulling identified strong theta during both behaviors; however, a marked reduction in theta frequency and the power of the theta harmonic were observed during pulling (**Figure 2A**, Welch PSD). Statistical analysis of frequency (**Figure 2B**) confirmed that theta frequency was reduced during string pulling (p = 0.0017, paired t-test). The power of the harmonic was also reduced during pulling relative to running (**Figure 2C**, p = 0.013, paired t-test).

**Figure 2.**
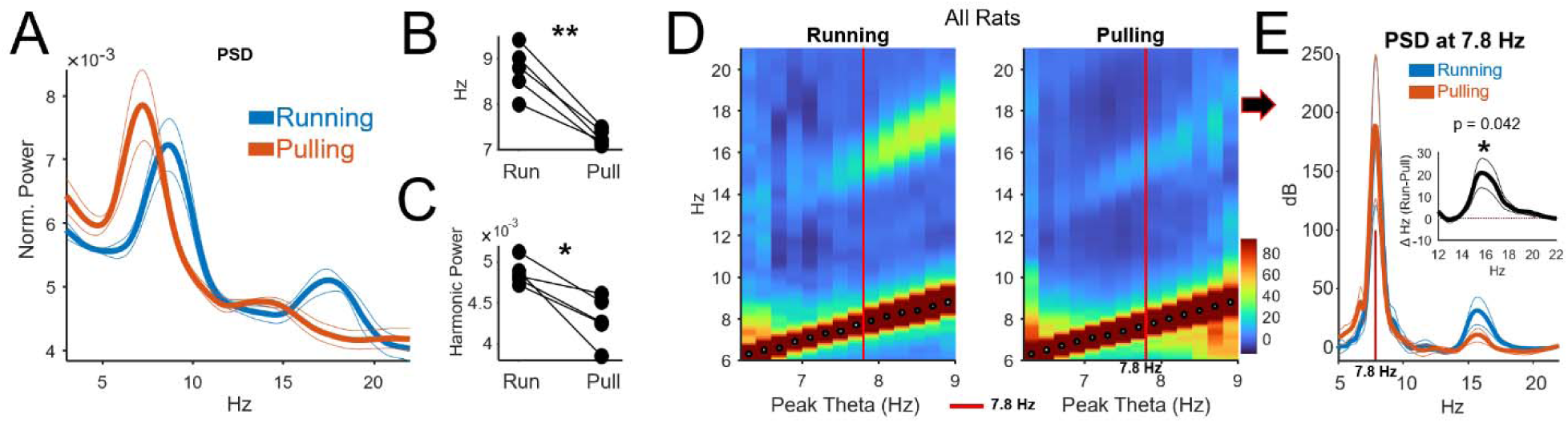
Slower theta and diminished harmonic during string pulling. A) Fourier Power-spectral density during track-running and string-pulling bouts for all 5 rats (Mean ±SEM). B) Theta frequency was reduced during string pulling relative to running in all animals (p = 0.0017, paired t-test). C) Harmonic power was also significantly reduced during string-pulling (p = 0.013, paired -test). D) The theta harmonic is a function of theta frequency during running. To determine if this is the case for string-pulling, the spectral response (Morelet Wavelet) was computed as a function of the frequency at the peak of the PSD at each speed (black and white dots). While there was a small indication of a harmonic during string pulling, it was reduced during relative to running. E) To ensure this effect held while controlling for theta frequency, we analyzed data at 7.8 Hz (vertical red lines in D and E). E shows the mean wavelet spectrogram at 7.8 Hz. The inset shows the within-animal difference (run-pull) averaged across animals mean ± SEM) showing a significantly larger harmonic during running when controlling for frequency. A paired t-test was performed to determine if the difference at 16 Hz was significant (p = 0.042).

The frequency of the harmonic is a function of the fundamental frequency and since the fundamental frequency of theta was lower during pulling, it was conceivable that this difference accounted for the reduced harmonic power. To control for this, harmonic power and the power-spectral density were computed for a range of theta frequencies during string-pulling and track running (**Figure 2D**). Visually, it appeared that the harmonic remained stronger during track running even when controlling for peak frequency. To test this, we selected a theta frequency that was clearly present during pulling and running (7.8 Hz, red vertical lines) and compared string and track-running PSDs at this frequency (**Figure 2E**). Analysis of the PSDs and the within-subject difference in the PSDs (**Figure 2E inset**) determined that power in the harmonic was reduced during string pulling when controlling for frequency (p = 0.042, paired t-test).

### The established positive relationship between movement speed and theta frequency is eliminated or reversed during string pulling

The positive relationships between theta frequency and power and running speed^20^ are believed to be essential for stabilizing place fields and the hippocampal spatial map considering naturally varying changes in movement speed. As with running towards a reward on a track, the string-pulling behavior, as implemented here, requires traveling a distance measured in the length of string pulled to acquire a reward. Thus, it was conceivable that the established relationship between string speed and theta frequency and power would be maintained during string pulling and potentially serve to stabilize neural representations of length pulled.

Consequently, we analyzed changes in the instantaneous theta frequency as a function of running and string speed. Instantaneous theta frequency was measured using the Hilbert transform (instfreq(), Matlab). An example from one recording session is presented in **Figure 3A**. To analyze frequency responses to speed across animals, the average wavelet spectrogram (n = 5 rats) was computed as a function of movement and string speed (**Figure 3B**). This figure illustrates the clear positive relationship between running speed and theta frequency. Notably, there was no positive relationship during string pulling. To increase the sensitivity of this analysis and to account for inter-animal differences in theta frequency and power, we normalized within-session measures of power and frequency such that they were between -1 (minimum) and +1 (maximum) (**Figure 3C**) and then averaged across animals (±SEM). These data indicate a clear dissociation between the relationship between running and pulling. As it is difficult to interpret standardized units, we also performed the within-session analysis using the original units (Hz) by subtracting the frequency observed at 10 cm/sec from all other measures (**Figure 3D**). Keeping the original units (Hz) demonstrated that the effect size of the speed-to-frequency relationship was notably larger (∼0.6 Hz) relative to string pulling (∼-0.1 Hz).

**Figure 3.**
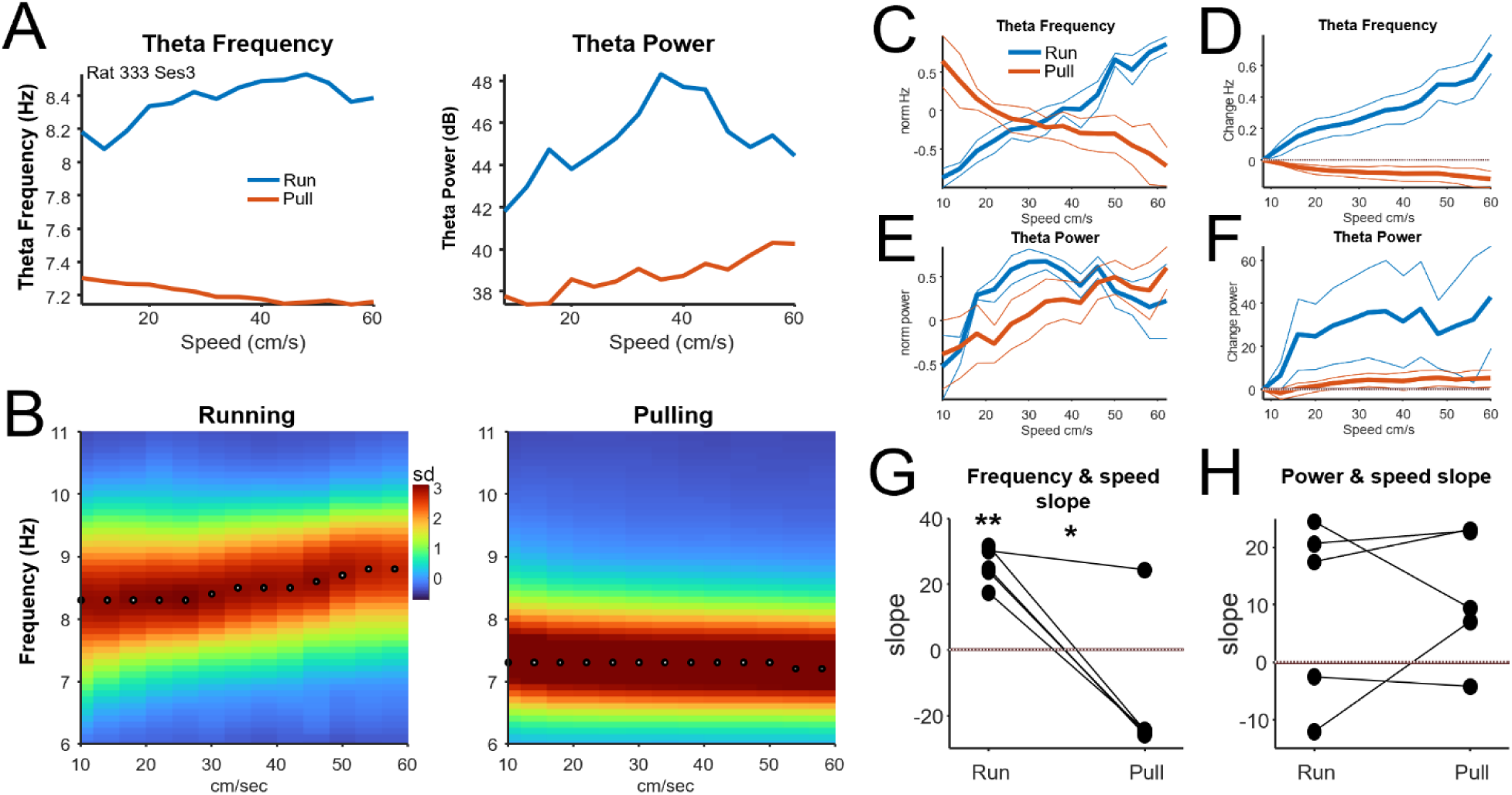
Theta frequency is not positively correlated with pulling speed. A) Theta frequency (left) and power (right) as a function of speed during running (blue) and string-pulling (orange) during a single recording session. B) Average wavelet spectrogram (n = 5 rats) computed for running (left) and pulling (right) speed (x axis). The expected positive relationship was observed during running, but not during string pulling. Dots indicate the peak of the power spectral density for each speed. Values are in units of standard deviation from the mean at each speed. E) Normalized theta frequency by speed (Mean ±SEM) where theta was standardized to -1 (minimum) and +1 (maximum) for each animal. While a clear positive relationship was observed between running and theta frequency, this relationship was reversed during string pulling. D) Same data as in C but where the data are Hz and normalized by subtracting the frequency value at the lowest speed (10 cm) from each individual value. This illustrates that the magnitude of the effect of speed on frequency during running was larger than during string pulling. E) As in C, but for the relationship between speed and power. F) As in D, but for power normalized to the power at the lowest speed. G) Slope of the relationship between speed and theta frequency for each animal during running and string pulling. There was a significant difference between string pulling and running (paired t-test, n = 5 rats, p = 0.011). While all slopes were positive during running, 4 of 5 were negative during string pulling. H) Slope of the relationship between speed and theta power during running and string pulling. There was no significant difference between running and string pulling (paired t-test, p = 0.732).

Statistical analysis of the slopes of the relationship between frequency and speed and power and speed are presented in **Figure 3GH**. Analysis of the slope between frequency and speed identified a significant difference between running and pulling (**Figure 3G**, p = 0.011, paired t-test, n = 5). All slopes were positive for track running (p = 0.001, t-test) and 4 of 5 slopes were negative during string pulling although the distribution did not differ statistically from zero (p = 0.2, t-test). In contrast, no difference between slopes for speed and theta power were observed (**Figure 3H**, p = 0.73, paired t-test). Notably, two of five animals did not exhibit a positive relationship between running speed and theta power, and visual inspection suggested that power rose but peaked at lower speeds than frequency (**Figure 3E**).

To summarize, we observed that the established positive relationship between theta frequency and movement speed does not hold for the goal-directed string-pulling behavior. The measure of string speed used here, however, integrates the motion of both paws and did not consider the reach-grasp cycle (**Figure 1B**). Consequently, we pursued a fine-grained analysis of speed and movement focused on the motion of the contralateral-to-implant (left) paw.

### Theta frequency peaks during the ‘push’ phase of the string-pulling behavior

To analyze the relationship between local-field activity and each phase of the reach-pull behavior, paw movement was segmented into lift, advance, grasp, pull, and push categories ^8^. Analysis of the speed of the left paw during each of these categories (**Figure 4A**) identified a strong effect of movement phase (p = 0.00001, F = 54.1, n = 5, within-subject ANOVA). The speed of paw movement was greatest during the lift and advance phases and lowest during the pull/push phases, presumably because push/pull required effort to advance the string forward. A similarly strong effect of acceleration by pull phase was also observed (**Figure 4B**, p = 0.00001, F = 61.9) with acceleration peaking during the lift phase.

**Figure 4.**
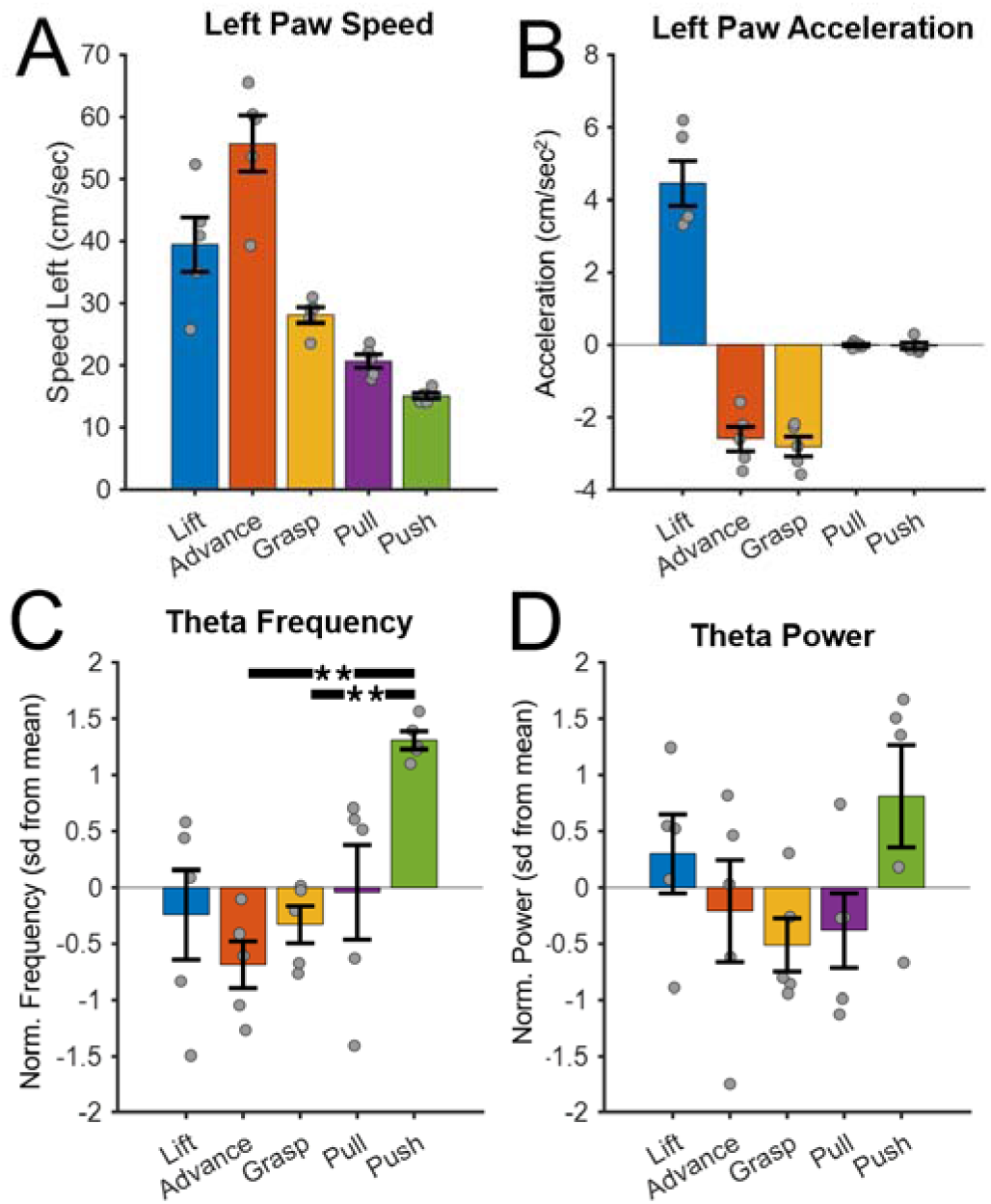
**Behavior and theta activity within each reach-pull cycle.** A) The speed of the left paw (Mean ±SEM) significantly varied by pull-phase (p = 0.00001, F = 54.1, n = 5, within-subject ANOVA). Each dot indicates a single rat. Speeds were higher during the lift and advance phases than the pull and push phases. B) Acceleration (cm/sec2) also varied significantly as a function of reach-pull phase (p = 0.00001, F = 61.9). C) Theta frequency peaked during the push phase. Y-axis indicates the z-score normalized theta power during each phase. A within ANOVA was performed (F = 7.692, p = 0.0012, within-subject ANOVA, post-hoc with Tukey-Kramer correction). D) No relationship between theta power and pull phase was observed (within subject ANOVA, F = 2.29, p = 0.11).

Instantaneous estimates of theta frequency were determined using the Hilbert method (instfreq(), Matlab) and averaged within each segmented phase of the reach-pull cycle for the left paw (**Figure 3C**). A strong main effect of pull phase was observed (p = 0.0012, F = 7.69, within-subject ANOVA) and post-hoc tests demonstrated that theta frequency peaked during the ‘push’ (p < 0.01, Tukey-Kramer correction). Interestingly, the push phase is the phase during which left paw movement was slowest.

Instantaneous estimates of theta power were measured as the absolute value of the Hilbert transform of the band-pass filtered theta signal (see Methods) and averaged within each segmented reach-pull phase (**Figure 4D**). Unlike theta frequency, there was no main effect of pull phase (within subject ANOVA, F = 2.29, p = 0.11).

### Theta becomes phase-locked during the ‘pull’ phase of the reach-pull cycle

To provide a more fine-grained analysis of the reach-pull behavior beyond the five categorized phases, the xy trajectories of the paw were transformed into degrees to produce a continuous circular measure of paw position (angle) at each phase of the reach-pull cycle. Paw speed as function of the circular coordinate frame is indicated in **Figure 5B** with degrees on the x-axis. The color bar at the bottom of the figure indicates the corresponding discrete pull-phase illustrated in **Figure 5A**. As illustrated in **Figure 5BC**, paw speed and acceleration at each phase of the reach-pull cycle was very consistent between animals, with speed peaking during the lift/advance phase and acceleration peaking at the beginning of the lift phase.

**Figure 5.**
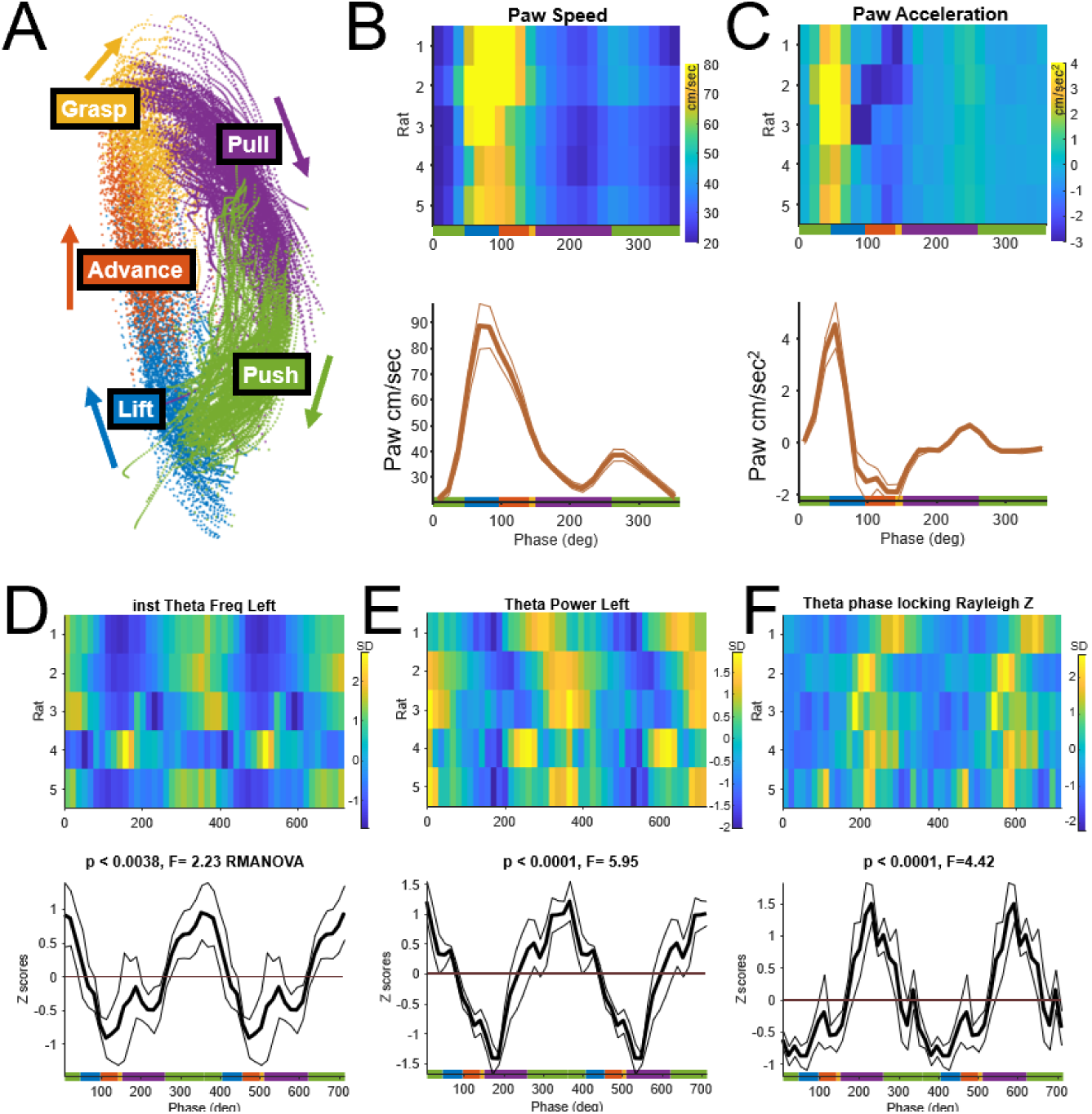
Theta features within each reach-pull cycle. A) Reach-pull trajectories from the left paw acquired from a single behavioral session color coded by the phase of the pull. B) Top plots the average speed of the left paw for each rat (row) with the mean ±SEM presented as the orange line in the subplot below. The dashed line indicates the speed of the string through each phase. The color-coded line at the bottom of the plot indicates the pull phase as illustrated in A. C) As in B, except for paw acceleration. D-E) Theta frequency, power, and theta phase locking to the reach-pull cycle. The top plots indicating theta features for each rat (row) and the bottom plot indicates the mean ±SEM. Two reach-pull cycles (720°) are presented on the x axis to improve visualization. Vertical dashed lines are to indicate the relationship between the peak of each feature and the reach-pull phase. D) Theta frequency changes within the reach-pull cycle for each rat (row) was normalized as z-scores to control for inter-animal differences in mean frequency. A clear effect of reach-pull phase was identified (within-subject ANOVA, p<0.001, F=3.03, eta2 =0.43) with frequency peaking during the ‘push’ phase. While significant, it should be noted that the absolute difference between the minimum and maximum frequency was small (0.1 Hz). E) As with frequency, a significant effect of reach-pull phase was observed for theta power (p<0.0001, F=5.25, eta2 =0.57) with frequency peaking during the ‘push’ phase. F) Phase locking to theta computed between each reach-pull cycle for each animal and measured as the Rayleigh z measure (circ_rtest). Significant theta phase locking across reach-pull cycles was observed (p<0.0001, F=4.42, eta2 =0.53) with phase locking peaking during the ‘pull’ phase.

Theta frequency and power were analyzed as a function of reach-pull cycle (**Figure 5D-E**). Consistent with the analysis presented in **Figure 4**, there was a strong effect of theta frequency (**Figure 5D**, p < 0.001, within-subject ANOVA, F = 3.03, eta^2^ = 0.43) with frequency peaking during the push phase and being lowest during the advance phase. A similar pattern was observed for theta power (**Figure 5E**, p < 0.0001, F = 5.25, eta^2^ = 0.57). This result differed from **Figure 4D** where no main effect of pull-phase was observed for power; however, it is conceivable that the additional precision offered by converting the pull phase into a continuous angular measure rather than discrete categories accounts for this discrepancy.

Theta phase locking indicates the extent the phase of the oscillation is consistent from trial to trial or behavior to behavior. Previous work suggests that theta phase locking peaks during meaningful instrumental behaviors such as pressing a button to receive reward or initiate movement^14,15^. Theta phase also aligns to each footstep when rats navigate a maze, but only when the animal is in a condition requiring spatial memory recall^18^. Thus, it was conceivable that phase of the theta oscillation aligns during reach-pull behaviors.

Theta phase locking to the reach-pull cycle was quantified using Rayleigh’s z statistic and plotted as a function of reach-pull phase (**Figure 5F**). We observed a significant effect of phase locking (p < 0.0001, within-subject ANOVA, F = 4.42, eta^2^ = 0.53) with the peak values occurring near the end of the pull cycle (right before the push). Interestingly, phase locking preceded the peak in theta frequency or power (**Figure 5DE**).

### String-pulling bouts end with a rapid and discrete increase in theta frequency

Visual inspection of the LFP suggested a notable increase in theta frequency at the end of each pulling bout. To further investigate, wavelet G+H) As in E+F but for period surrounding the end of each pulling bout. At the end of a bout, there was a rapid increase in theta frequency of ∼1Hz as indicated in H (p = 0.00093, paired t-test).

spectrograms were generated and aligned to the start and end of each pull bout. For example, **Figure 6AB** shows data from a single animal/session. The speed of the rat’s nose is overlaid (white line) on each spectrogram and string speed is indicated in **Figure 6CD**. As shown in **Figure 6B**, there was a rapid ∼1 Hz increase in theta frequency aligned precisely to the end of each bout. This increase also aligned with the rapid increase in nose speed occurring when the rat turned around to retrieve the food reward. Thus, the rapid increase in theta frequency could result from the large increase in vestibular drive accompanying the turn.

**Figure 6.**
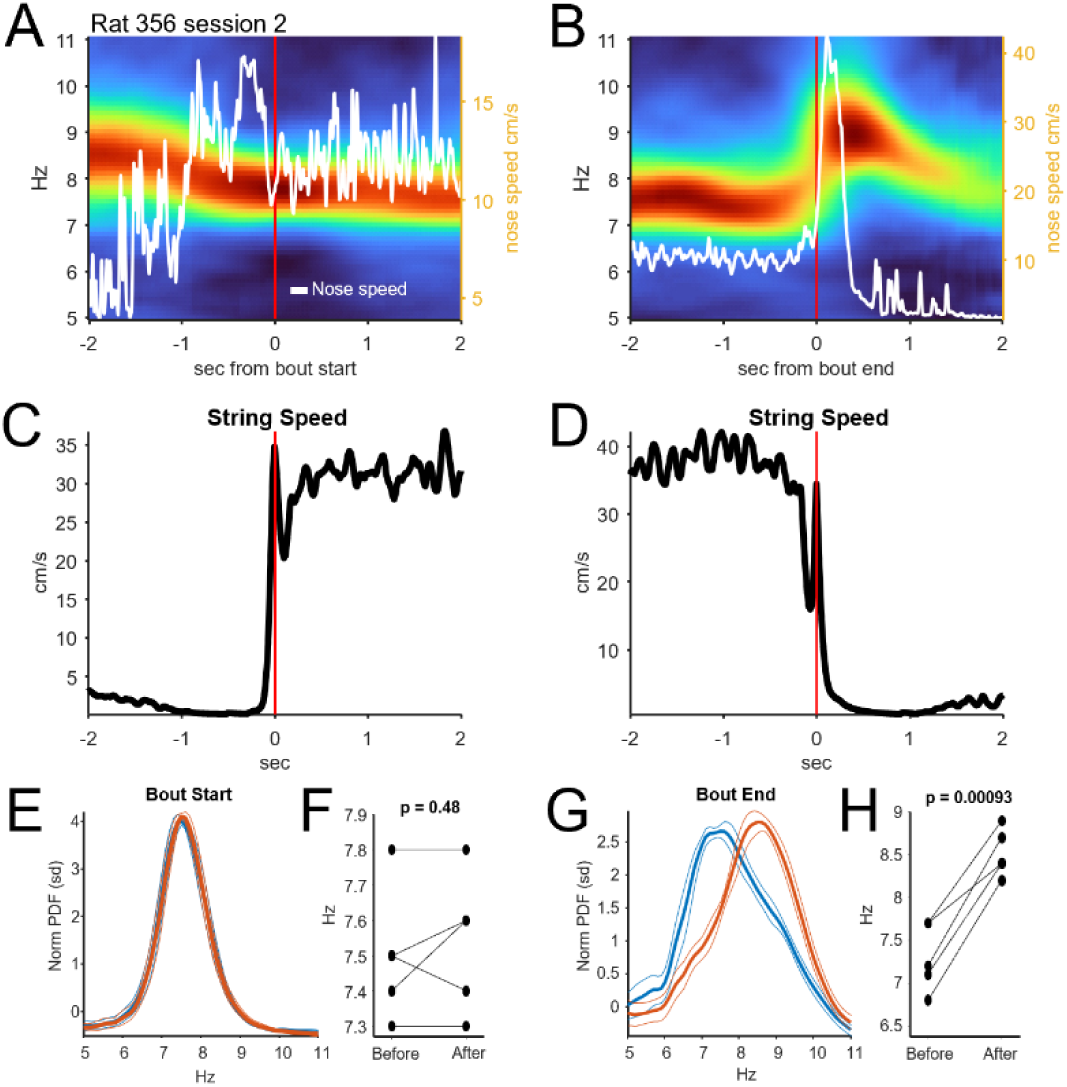
Theta-band responses to the start and end of each string-pulling bout. A) Example of the mean wavelet spectrogram and nose speed (white line) aligned to the start of the pulling bouts during a single session. Pulling bout onset was determined by analyzing string movement. B) As in A but aligned to bout end. Note the large increase in theta that occurred simultaneously with nose movement. The nose moved as the rat turned around to retrieve food reward at the end of the bout. C+D) Mean string speed aligned to the start and end of the pulling bout. E) To quantify changes in frequency as a function of bout start and end, the PSD for the period before (-250 ms to -50 ms) and after (+50 ms to +250 ms) bout start was measured for each animal. The mean normalized PSD for before (blue) and after (orange) is presented in E. F) As suggested in E, there was no difference in theta frequency before and after bout start (p = 0.48, paired t-test).

To determine if this effect was consistent across animals, the wavelet PSDs for the time intervals preceding (-250 to -50 ms) and following (+50 to +250 ms) bout onset and offset were computed for each rat and then averaged (**Figure 6EG**). No change in theta frequency was observed surrounding bout onset (p = 0.48, paired t-test); however, a significant >1 Hz increase in theta frequency was observed at bout end (p = 0.00093, **Figure 6H**).

## Discussion

Hippocampal neuronal firing rates^21^, theta oscillation power, and theta frequency^2,22^ rise with increasing running speed. These adaptive responses may be critical for stabilizing neural representations of position in the face of varying motion^1,5^, which, in turn, may contribute to episodic memory^5^. Long-term potentiation (LTP) is also modulated by the phase at which neurons fire relative to theta, suggesting a role for theta in guiding synaptic changes required for memory formation^3,4^. Modulation by phase also suggests that salient inputs to the hippocampus must be precisely timed relative to theta oscillation.

Real-time estimation of movement speed could be derived from a variety of hippocampal inputs including visual flow, vestibular signals, physical effort, and proprioceptive self-motion. There is conflicting evidence regarding which factors or combinations thereof drive or enable the frequency-speed relationship. For example, a positive relationship between theta frequency and speed has been reported in head-fixed guinea pigs walking on a treadmill (speeds < 6 cm/sec)^22^ and untethered dogs running on treadmills^23^. This suggests that sensorimotor information related to limb movement is sufficient to change theta frequency. These data, however, conflict with some studies involving treadmill-running rats^20,24^ that found no relationship between theta frequency and speed, suggesting that sensorimotor input from leg/arm motion is not sufficient.

Adding visual flow information appears to restore the theta-speed relationship as theta frequency is positively correlated with speed in head-fixed mice running in virtual reality environments^25^, although the effect size in virtual reality environments is notably reduced relative to real-world exploration.

In these studies, and in most investigations of spatial processing in rodents, position is tracked as the location of the head or the center of mass of the body. Consequently, little is known about how theta synchronizes to fine motor movements, such as forelimb movements. Studies that have investigated limb end eye motion have observed that theta can synchronize to button-presses^14^, eye saccades^15^, and footsteps^18^, but mostly when such actions relate to memory encoding or retrieval. We predicted that theta activity would also synchronize to body movement in a non-spatial task with minimal memory requirements, but where actions (pull/push of a string) propelled the animals forward in the dimension of ‘string length’ towards a food reward^11^. Despite this task placing multiple speed-associated cues into conflict (e.g., stationary hindlimb, moving forelimb, minimal flow field or vestibular signaling) we found that this behavior induced robust hippocampal theta (**Figure 2A**) and allowed for detailed analysis of each reach/pull movement.

### Theta frequency is notably reduced during string pulling relative to running

We found that theta frequency was consistently lower during string-pulling relative to track-running. Indeed, theta frequency at the fastest pulling speeds (60 cm/sec, theta: ∼7.2 Hz) was lower than at the slowest running speeds (10 cm/sec, theta: ∼8.2 Hz, see **Figure 3B**).

Diminished theta frequency has also been reported in motor tasks with reduced sensorimotor or vestibular drive^26,27^ including virtual reality navigation^28–31^. The study by Safaryan & Mehta^30^ was unique in that they observed that while theta frequency was reduced in virtual reality, theta power increased, a factor that could reflect the impact of strong visual input on hippocampal processing. Taken together, reduced theta frequency observed during string pulling likely results from reduced visual, vestibular, and sensorimotor drive. That said, this ideas does correspond well to observations by Czurkó et al.^24^ who did not observe any difference in peak frequency between treadmill running and running/walking, despite the notable reduction in flow-field and vestibular information during treadmill running. This suggests that the elimination of hindlimb movement during string pulling was sufficient to significantly reduce theta relative to treadmill walking.

### Reduced theta harmonic during string pulling

In addition to the theta frequency, the theta harmonic^32^ was also reduced during string-pulling relative to track running, even after controlling for attenuation due to reduced theta frequency (**Figure 2**). There is debate regarding the physiological origins of the harmonic as harmonics can be produced by multiple interacting mechanisms^33^. For example, they can result from higher-frequency oscillations being phase-locked to theta^34^, or to asymmetries in the shape of an oscillatory waveform^31,35–38^. Asymmetries, for example, could result from subpopulations of interneurons and principal cells firing at distinct phases of theta^31^. The notable variety of interneuron subtypes expressing distinct tuning for phases of theta and gamma^39,40^ contribute to asymmetries in extracellular currents during a theta cycle. Given that hippocampal inputs convey vestibular, visual, and somatosensory input to distinct neural subpopulations^29,41,42^, it is conceivable that the reduced harmonic power observed during string pulling is a consequence of fewer of these subgroups being activated.

### The theta-frequency-to-speed relationship is absent or reversed during string-pulling

We found that during string pulling the established positive relationship between movement speed and theta frequency is not present. Indeed, gross measures of string speed (**Figure 3**) were often negatively correlated with theta frequency. This observation parallels results from experiments in rats running on treadmills that also failed to identify a relationship between theta frequency and speed^20,24^, although the Czurkó study did observe that individual CA1 neurons increased firing with increased speed, suggesting that speed-related input is still drive neural activity in the treadmill behavior. More recent work suggest that adding visual flow information to the task, such as through virtual reality, is sufficient to restore the theta-speed relationship^25^. Thus, forepaw movement, even in a task that required animals to move in ‘string space’ was not sufficient to generate the speed-theta frequency relationship. From a theoretical perspective, the need to maintain place-field stability despite varying speeds in real or virtual 2D spaces is clear. It is less clear whether such stability is required when moving through ‘string space’, at least under the training conditions used here.

### Theta frequency and power peak during the push phase of string pulling

Moving from gross measures of movement and string speed to fine-grained measures of forepaw movement yielded further insights into the speed-theta relationship. Data in **Figures 4** and **5** indicate that theta power and frequency peaked during the ‘push’, when contralateral paw speed was at its minimum (**Figure 5DE**). Relative to the lift and advance phases, the pushing movement is also the most effortful and the action most directly connected with advancing the string to the goal. The pull/push phase shares similarities with the ‘push’ of the paw backwards during walking or running which is also the phase that most directly advances the animal through space. This suggests that, during running, theta frequency and power will peak during the backward push of the contralateral limb once it contacts the floor surface. To our knowledge, this question has not yet been investigated. That said, analysis of such behavior is complicated by the fact that running involves four limbs with interdependent phase-locked movements. Thus, theta could respond to higher-order features of inter-limb interactions.

An alternative to the above explanation is that theta frequency is modulated by effort. The brief increase in theta frequency during the push could result from the physical effort required to pull and push the string forward. While it is difficult to dissociate effort from movement speed in most behaviors, previous studies have reported increased theta frequency and power during jumping where both theta frequency and power increased with the distance of an anticipated jump^12,13^.

### Theta oscillations phase-lock during the pull phase of string-pulling

Synchronization between oscillatory activity and behavior has been observed in sensorimotor regions of cortex^43^ and in higher order associative regions such as the hippocampus^18^. In sensory and motor cortex, such synchrony likely facilitates active sensing^43,44^ and the attentional gating of sensory information^45^. The function of body-oscillatory synchrony in higher-order association regions such as the hippocampus is less clear, although recent data suggests that synchrony may also facilitate the gating of sensorimotor information into the hippocampus^16–18^. Beyond gating, theta phase-locking may also support inter-regional communication^46–51^.

Previous studies have reported that the phase of hippocampal theta aligns to meaningful actions, such as button presses or eye saccades that predicted future reinforcement^14,15^ or to individual footsteps, but only when memory demands are high^18^. The extent to which a ‘meaningful’ action or cognitive process modulates phase-coupling may be a matter of degree. For instance, one study with head-fixed mice running on a ball found weak theta-to-stepping phase-locking in a task with limited cognitive demands^52^. To our knowledge, no study to date has performed a fine-grained analysis of theta phase-locking to movements involved in ‘lifting, ‘advancing’, ‘grasping’, ‘pushing’, and ‘pulling’. Using the string-pulling task, we observed that the phase of the theta oscillation aligned to the pull phase (**Figure 5F**). The pull is the first phase where the animal’s movement directly and causally advances the string to receive the reward.

This finding is in agreement with other observations indicating that theta phase aligns or “resets” to task-meaningful actions such as a lever-pressing for brain stimulation^53^ or food reinforcement^54^. Other rhythms may also synchronize to the pull phase given evidence for coupling between hippocampal theta and respiratory and whisking movements^16,17^. Theta phase alignment during the pull phase may also reflect sensorimotor processing and gating. The pull phase immediately follows the ‘grasp’ and likely involves considerable tactile feedback in response to grasping and pulling string. Indeed, work by Grion et al.^16^ indicated that coherence between hippocampal theta and whisking movements increases during texture discrimination.

They also observed that rats were more successful at texture identification when theta-to-whisking coherence was high, suggesting that sensorimotor information from the whiskers was more effectively transmitted to higher-order centers during phase locking. For the string-pulling behavior reported here, the approach to the reward is measured in string length. Given that the pull and push are the phases most linked to progression to the goal, it is conceivable that gating could facilitate hippocampal computations of the progress in string-length towards the goal.

### Large discrete increase in theta frequency at the end of each pulling bout

At the end of each pulling bout, the rats rapidly turned 180° to approach the food reward spout. At this moment, we observed a strong and step-like increase in theta frequency that aligned precisely to head turning (**Figure 6B**). The magnitude of this effect was large (∼1.5 Hz), likely resulting from the rapid increase in vestibular drive produced by head rotation.

Considerable evidence indicates a role of vestibular input in driving theta-band activity in the hippocampus^55^. Increased theta frequency may also be associated with a neural “shift” or state change while transitioning toward a new expected outcome such as a different reward^56,57^.

### Limitations

This study used fixed arrays of wire tetrodes. These arrays were made by hand and thus lacked the precise inter-electrode spacing and density of silicon probes. As a result, we were not able to perform current-source density analysis to unambiguously determine the depth of each probe by hippocampal layer. The polarity, power, and asymmetry of theta can vary by depth^58,59^ and so we focused on within-animal relative changes in power and refrained from analysis of asymmetry. Planned studies using high-density silicon probes will permit a full analysis of asymmetry.

Another limitation is that these data do not allow us to definitively distinguish between Type I and Type II theta^60^. Type I theta peaks during movement, has a higher frequency than Type II theta, and is not affected by cholinergic antagonists (e.g., atropine). In contrast, Type II theta is more involved in awake immobility and is atropine sensitive, although it can be triggered by sensory stimuli^61^. Thus, theta activity measured here, as in many studies, is potentially a superposition of Type I and Type II theta.

While we focused on the contralateral paw, anatomical evidence indicates that the hippocampus and the lateral entorhinal cortex, a major cortical input, receive somatosensory information from both sides of the body^62,63^. The hippocampus also has extensive commissural projections^64^. Furthermore, the string-pulling behavior required notable inter-limb coordination to guide the string from one paw to the next, suggesting significant between-hemispheric interactions. Thus, we cannot definitively conclude that theta only synchronized to the contralateral paw. Future studies comparing single limb vs. bimanual behaviors could help determine the relative functional contribution of each limb to theta-action.

## Conclusions

In the task used here, rats learned to pull a fixed length of string for food reward. Since reward delivery was purely a function of the animal’s self-initiated movement through ‘string space’ towards a fixed length of string, we predicted that established relationships between theta frequency and running would also apply to the string-pulling behavior. Given that the speed-frequency relationship has been reported in virtual reality environments, we felt this relationship would hold despite the lack of vestibular and hindlimb self-motion signals. Contrary to this prediction, we observed that relationships between theta oscillations and movement were distinctly different during running versus string-pulling. While theta remained robust in both conditions, frequency was significantly reduced during string-pulling and showed no speed-frequency coupling, unlike running. This difference could result from the conflict between hindlimb and forelimb signals, with only forelimb movement signaling advancement. Fine-grained analysis of paw movements during string-pulling (lift, advance, grasp, pull, push) revealed that theta power and frequency peaked during the contralateral paw’s downward push despite paw speed being low during this action. This is consistent with the idea that theta can respond to effort rather than purely kinematic information. Notably, running-associated theta may similarly reflect both speed and effort signals as most locomotor tasks conflate these variables. Finally, theta phase preferentially aligned to the downward pull of the string, the first action that directly advances the string forward. Since phase-locking has been associated with sensorimotor gating, synchrony at this point could reflect the gating of inputs that are the most causally relevant for reaching the reward, potentially facilitating integration of causal action-outcome signals for memory encoding and navigation. Taken together, these data support a dual-scale view of hippocampal processing and theta-band activity where macroscale theta activity requires suprathreshold visual, vestibular, and sensorimotor drive and microscale theta remains sensitive to subsecond limb movements.

## Acknowledgments

Funding:

McKnight Brain Research Foundation NIH R01 RF1AG081767

R01 NIH BRAIN NS123424-01

Arizona Alzheimer’s Consortium Pilot Grant

## References

1. Geisler, C., Robbe, D., Zugaro, M., Sirota, A., and Buzsáki, G. (2007). Hippocampal place cell assemblies are speed-controlled oscillators. Proc. Natl. Acad. Sci. U. S. A. 104, 8149– 8154. 10.1073/pnas.0610121104.

2. McFarland, W.L., Teitelbaum, H., and Hedges, E.K. (1975). Relationship between hippocampal theta activity and running speed in the rat. J. Comp. Physiol. Psychol. 88, 324– 328.

3. Hölscher, C., Anwyl, R., and Rowan, M.J. (1997). Stimulation on the positive phase of hippocampal theta rhythm induces long-term potentiation that can Be depotentiated by stimulation on the negative phase in area CA1 in vivo. J. Neurosci. Off. J. Soc. Neurosci. 17, 6470–6477. 10.1523/JNEUROSCI.17-16-06470.1997.

4. Huerta, P.T., and Lisman, J.E. (1993). Heightened synaptic plasticity of hippocampal CA1 neurons during a cholinergically induced rhythmic state. Nature 364, 723–725. 10.1038/364723a0.

5. O’Keefe, J., and Nadel, L. (1978). The Hippocampus as a Cognitive Map (Clarendon Press).

6. Jacobs, I.F., and Osvath, M. (2015). The string-pulling paradigm in comparative psychology. J. Comp. Psychol. 129, 89–120. 10.1037/a0038746.

7. Singh, S., Mandziak, A., Barr, K., Blackwell, A.A., Mohajerani, M.H., Wallace, D.G., and Whishaw, I.Q. (2019). Human string-pulling with and without a string: movement, sensory control, and memory. Exp. Brain Res. 237, 3431–3447. 10.1007/s00221-019-05684-y.

8. Blackwell, A.A., Banovetz, M.T., Qandeel, Whishaw, I.Q., and Wallace, D.G. (2018). The structure of arm and hand movements in a spontaneous and food rewarded on-line string-pulling task by the mouse. Behav. Brain Res. 345, 49–58. 10.1016/j.bbr.2018.02.025.

9. Blackwell, A.A., Köppen, J.R., Whishaw, I.Q., and Wallace, D.G. (2018). String-pulling for food by the rat: Assessment of movement, topography and kinematics of a bilaterally skilled forelimb act. Learn. Motiv. 61, 63–73. 10.1016/j.lmot.2017.03.010.

10. Blackwell, A.A., Widick, W.L., Cheatwood, J.L., Whishaw, I.Q., and Wallace, D.G. (2018). Unilateral forelimb sensorimotor cortex devascularization disrupts the topographic and kinematic characteristics of hand movements while string-pulling for food in the rat. Behav. Brain Res. 338, 88–100. 10.1016/j.bbr.2017.10.014.

11. Jordan, G.A., Vishwanath, A., Holguin, G., Bartlett, M.J., Tapia, A.K., Winter, G.M., Sexauer, M.R., Stopera, C.J., Falk, T., and Cowen, S.L. (2024). Automated system for training and assessing reaching and grasping behaviors in rodents. J. Neurosci. Methods 401, 109990. 10.1016/j.jneumeth.2023.109990.

12. Vanderwolf, C.H. (1969). Hippocampal electrical activity and voluntary movement in the rat. Electroencephalogr Clin Neurophysiol 26, 407–418.

13. Bland, B.H., Jackson, J., Derrie-Gillespie, D., Azad, T., Rickhi, A., and Abriam, J. (2006). Amplitude, frequency, and phase analysis of hippocampal theta during sensorimotor processing in a jump avoidance task. Hippocampus 16, 673–681. 10.1002/hipo.20210.

14. Ter Wal, M., Linde-Domingo, J., Lifanov, J., Roux, F., Kolibius, L.D., Gollwitzer, S., Lang, J., Hamer, H., Rollings, D., Sawlani, V., et al. (2021). Theta rhythmicity governs human behavior and hippocampal signals during memory-dependent tasks. Nat. Commun. 12, 7048. 10.1038/s41467-021-27323-3.

15. Jutras, M.J., Fries, P., and Buffalo, E.A. (2013). Oscillatory activity in the monkey hippocampus during visual exploration and memory formation. Proc. Natl. Acad. Sci. U. S. A. 110, 13144–13149. 10.1073/pnas.1302351110.

16. Grion, N., Akrami, A., Zuo, Y., Stella, F., and Diamond, M.E. (2016). Coherence between Rat Sensorimotor System and Hippocampus Is Enhanced during Tactile Discrimination. PLoS Biol. 14, e1002384. 10.1371/journal.pbio.1002384.

17. Kleinfeld, D., Deschênes, M., and Ulanovsky, N. (2016). Whisking, Sniffing, and the Hippocampal θ-Rhythm: A Tale of Two Oscillators. PLoS Biol. 14, e1002385. 10.1371/journal.pbio.1002385.

18. Joshi, A., Denovellis, E.L., Mankili, A., Meneksedag, Y., Davidson, T.J., Gillespie, A.K., Guidera, J.A., Roumis, D., and Frank, L.M. (2023). Dynamic synchronization between hippocampal representations and stepping. Nature, 1–7. 10.1038/s41586-023-05928-6.

19. Mathis, A., Mamidanna, P., Cury, K.M., Abe, T., Murthy, V.N., Mathis, M.W., and Bethge, M. (2018). DeepLabCut: markerless pose estimation of user-defined body parts with deep learning. Nat. Neurosci. 21, 1281–1289. 10.1038/s41593-018-0209-y.

20. Whishaw, I.Q., and Vanderwolf, C.H. (1973). Hippocampal EEG and behavior: changes in amplitude and frequency of RSA (theta rhythm) associated with spontaneous and learned movement patterns in rats and cats. Behav. Biol. 8, 461–484. 10.1016/s0091-6773(73)80041-0.

21. McNaughton, B.L., Barnes, C.A., and O’Keefe, J. (1984). The contributions of position, direction, and velocity to single unit activity in the hippocampus of freely-moving rats. Exp. Brain Res. 54, 195. 10.1007/BF00235832.

22. Rivas, J., Gaztelu, J.M., and García-Austt, E. (1996). Changes in hippocampal cell discharge patterns and theta rhythm spectral properties as a function of walking velocity in the guinea pig. Exp. Brain Res. 108, 113–118. 10.1007/BF00242908.

23. Arnolds, D.E., Lopes da Silva, F.H., Aitink, J.W., and Kamp, A. (1979). Hippocampal EEG and behaviour in dog. II. Hippocampal EEG correlates with elementary motor acts. Electroencephalogr. Clin. Neurophysiol. 46, 571–580. 10.1016/0013-4694(79)90010-5.

24. Czurkó, A., Hirase, H., Csicsvari, J., and Buzsáki, G. (1999). Sustained activation of hippocampal pyramidal cells by “space clamping” in a running wheel. Eur. J. Neurosci. 11, 344–352. 10.1046/j.1460-9568.1999.00446.x.

25. Chen, G., King, J.A., Lu, Y., Cacucci, F., and Burgess, N. (2018). Spatial cell firing during virtual navigation of open arenas by head-restrained mice. eLife 7, e34789. 10.7554/eLife.34789.

26. Bland, B.H., and Oddie, S.D. (2001). Theta band oscillation and synchrony in the hippocampal formation and associated structures: the case for its role in sensorimotor integration. Behav. Brain Res. 127, 119–136. 10.1016/S0166-4328(01)00358-8.

27. Foster, T.C., Castro, C.A., and McNaughton, B.L. (1989). Spatial selectivity of rat hippocampal neurons: dependence on preparedness for movement. Science 244, 1580–1582. 10.1126/science.2740902.

28. Ekstrom, A.D., Caplan, J.B., Ho, E., Shattuck, K., Fried, I., and Kahana, M.J. (2005). Human hippocampal theta activity during virtual navigation. Hippocampus 15, 881–889. 10.1002/hipo.20109.

29. Ravassard, P., Kees, A., Willers, B., Ho, D., Aharoni, D.A., Cushman, J., Aghajan, Z.M., and Mehta, M.R. (2013). Multisensory control of hippocampal spatiotemporal selectivity. Science 340, 1342–1346. 10.1126/science.1232655.

30. Safaryan, K., and Mehta, M.R. (2021). Enhanced hippocampal theta rhythmicity and emergence of eta oscillation in virtual reality. Nat. Neurosci. 24, 1065–1070. 10.1038/s41593-021-00871-z.

31. Terrazas, A., Krause, M., Lipa, P., Gothard, K.M., Barnes, C.A., and McNaughton, B.L. (2005). Self-motion and the hippocampal spatial metric. J. Neurosci. Off. J. Soc. Neurosci. 25, 8085–8096. 10.1523/JNEUROSCI.0693-05.2005.

32. Harper, R.M. (1971). Frequency changes in hippocampal electrical activity during movement and tonic immobility. Physiol. Behav. 7, 55–58. 10.1016/0031-9384(71)90235-6.

33. Dellavale, D., Velarde, O.M., Mato, G., and Urdapilleta, E. (2020). Complex interplay between spectral harmonicity and different types of cross-frequency couplings in nonlinear oscillators and biologically plausible neural network models. Phys. Rev. E 102, 062401. 10.1103/PhysRevE.102.062401.

34. Scheffer-Teixeira, R., Belchior, H., Leao, R.N., Ribeiro, S., and Tort, A.B.L. (2013). On High-Frequency Field Oscillations (>100 Hz) and the Spectral Leakage of Spiking Activity. J. Neurosci. 33, 1535–1539. 10.1523/JNEUROSCI.4217-12.2013.

35. Buzsáki, G., Czopf, J., Kondákor, I., and Kellényi, L. (1986). Laminar distribution of hippocampal rhythmic slow activity (RSA) in the behaving rat: current-source density analysis, effects of urethane and atropine. Brain Res. 365, 125–137. 10.1016/0006-8993(86)90729-8.

36. Cole, S.R., and Voytek, B. (2017). Brain Oscillations and the Importance of Waveform Shape. Trends Cogn. Sci. 21, 137–149. 10.1016/j.tics.2016.12.008.

37. Kennedy, J.P., Zhou, Y., Qin, Y., Lovett, S.D., Sheremet, A., Burke, S.N., and Maurer, A.P. (2022). A Direct Comparison of Theta Power and Frequency to Speed and Acceleration. J. Neurosci. 42, 4326–4341. 10.1523/JNEUROSCI.0987-21.2022.

38. Sheremet, A., Burke, S.N., and Maurer, A.P. (2016). Movement enhances the nonlinearity of hippocampal theta. J. Neurosci. 36, 4218–4230. 10.1523/JNEUROSCI.3564-15.2016.

39. Freund, T.F., and Buzsáki, G. (1996). Interneurons of the hippocampus. Hippocampus 6, 347–470. 10.1002/(SICI)1098-1063(1996)6:4<347::AID-HIPO1>3.0.CO;2-I.

40. Pelkey, K.A., Chittajallu, R., Craig, M.T., Tricoire, L., Wester, J.C., and McBain, C.J. (2017). Hippocampal GABAergic Inhibitory Interneurons. Physiol. Rev. 97, 1619–1747. 10.1152/physrev.00007.2017.

41. Hüfner, K., Strupp, M., Smith, P., Brandt, T., and Jahn, K. (2011). Spatial separation of visual and vestibular processing in the human hippocampal formation. Ann. N. Y. Acad. Sci. 1233, 177–186. 10.1111/j.1749-6632.2011.06115.x.

42. Inayat, S., McAllister, B.B., Whishaw, I.Q., and Mohajerani, M.H. (2023). Hippocampal conjunctive and complementary CA1 populations relate sensory events to movement. iScience 26. 10.1016/j.isci.2023.106481.

43. Leszczynski, M., and Schroeder, C.E. (2019). The Role of Neuronal Oscillations in Visual Active Sensing. Front. Integr. Neurosci. 13, 32. 10.3389/fnint.2019.00032.

44. Ahrens, K.F., and Kleinfeld, D. (2004). Current flow in vibrissa motor cortex can phase-lock with exploratory rhythmic whisking in rat. J. Neurophysiol. 92, 1700–1707. 10.1152/jn.00020.2004.

45. Lakatos, P., Karmos, G., Mehta, A.D., Ulbert, I., and Schroeder, C.E. (2008). Entrainment of Neuronal Oscillations as a Mechanism of Attentional Selection. Science 320, 110–113. 10.1126/science.1154735.

46. Benchenane, K., Peyrache, A., Khamassi, M., Tierney, P.L., Gioanni, Y., Battaglia, F.P., and Wiener, S.I. (2010). Coherent theta oscillations and reorganization of spike timing in the hippocampal-prefrontal network upon learning. Neuron 66, 921–936. 10.1016/j.neuron.2010.05.013.

47. Hyman, J.M., Zilli, E.A., Paley, A.M., and Hasselmo, M.E. (2010). Working Memory Performance Correlates with Prefrontal-Hippocampal Theta Interactions but not with Prefrontal Neuron Firing Rates. Front. Integr. Neurosci. 4, 2. 10.3389/neuro.07.002.2010.

48. Jones, M.W., and Wilson, M.A. (2005). Theta rhythms coordinate hippocampal-prefrontal interactions in a spatial memory task. PLoS Biol. 3, e402. 10.1371/journal.pbio.0030402.

49. Siapas, A.G., Lubenov, E.V., and Wilson, M.A. (2005). Prefrontal phase locking to hippocampal theta oscillations. Neuron 46, 141–151. 10.1016/j.neuron.2005.02.028.

50. Sigurdsson, T., Stark, K.L., Karayiorgou, M., Gogos, J.A., and Gordon, J.A. (2010). Impaired hippocampal-prefrontal synchrony in a genetic mouse model of schizophrenia. Nature 464, 763–767. 10.1038/nature08855.

51. Sirota, A., Montgomery, S., Fujisawa, S., Isomura, Y., Zugaro, M., and Buzsáki, G. (2008). Entrainment of neocortical neurons and gamma oscillations by the hippocampal theta rhythm. Neuron 60, 683–697. 10.1016/j.neuron.2008.09.014.

52. Joshi, A., and Somogyi, P. (2020). Changing phase relationship of the stepping rhythm to neuronal oscillatory theta activity in the septo-hippocampal network of mice. Brain Struct. Funct. 225, 871–879. 10.1007/s00429-020-02031-8.

53. Buño, W., and Velluti, J.C. (1977). Relationships of hippocampal theta cycles with bar pressing during self-stimulation. Physiol. Behav. 19, 615–621. 10.1016/0031-9384(77)90035-x.

54. Semba, K., and Komisaruk, B.R. (1978). Phase of the theta wave in relation to different limb movements in awake rats. Electroencephalogr. Clin. Neurophysiol. 44, 61–71. 10.1016/0013-4694(78)90105-0.

55. Aitken, P., Zheng, Y., and Smith, P.F. (2018). The modulation of hippocampal theta rhythm by the vestibular system. J. Neurophysiol. 119, 548–562. 10.1152/jn.00548.2017.

56. Jamali, S., Dezfouli, M.P., Kalbasi, A., Daliri, M.R., and Haghparast, A. (2023). Selective Modulation of Hippocampal Theta Oscillations in Response to Morphine versus Natural Reward. Brain Sci. 13, 322. 10.3390/brainsci13020322.

57. Lansink, C.S., Meijer, G.T., Lankelma, J.V., Vinck, M.A., Jackson, J.C., and Pennartz, C.M.A. (2016). Reward Expectancy Strengthens CA1 Theta and Beta Band Synchronization and Hippocampal-Ventral Striatal Coupling. J. Neurosci. 36, 10598–10610. 10.1523/JNEUROSCI.0682-16.2016.

58. Montgomery, S.M., Sirota, A., and Buzsáki, G. (2008). Theta and gamma coordination of hippocampal networks during waking and rapid eye movement sleep. J. Neurosci. 28, 6731– 6741. 10.1523/JNEUROSCI.1227-08.2008.

59. Buzsáki, G., Rappelsberger, P., and Kellényi, L. (1985). Depth profiles of hippocampal rhythmic slow activity (’theta rhythm’) depend on behaviour. Electroencephalogr. Clin. Neurophysiol. 61, 77–88. 10.1016/0013-4694(85)91075-2.

60. Kramis, R., Vanderwolf, C.H., and Bland, B.H. (1975). Two types of hippocampal rhythmical slow activity in both the rabbit and the rat: relations to behavior and effects of atropine, diethyl ether, urethane, and pentobarbital. Exp. Neurol. 49, 58–85. 10.1016/0014-4886(75)90195-8.

61. Sainsbury, R.S., Heynen, A., and Montoya, C.P. (1987). Behavioral correlates of hippocampal type 2 theta in the rat. Physiol. Behav. 39, 513–519. 10.1016/0031-9384(87)90382-9.

62. Burwell, R.D., and Amaral, D.G. (1998). Cortical afferents of the perirhinal, postrhinal, and entorhinal cortices of the rat. J Comp Neurol 398, 179–205.

63. Canto, C.B., Wouterlood, F.G., and Witter, M.P. (2008). What Does the Anatomical Organization of the Entorhinal Cortex Tell Us? Neural Plast. 2008, 381243. 10.1155/2008/381243.

64. Demeter, S., Rosene, D.L., and Van Hoesen, G.W. (1985). Interhemispheric pathways of the hippocampal formation, presubiculum, and entorhinal and posterior parahippocampal cortices in the rhesus monkey: the structure and organization of the hippocampal commissures. J. Comp. Neurol. 233, 30–47. 10.1002/cne.902330104.

