## Supplementary Figures for "Opposing responses of hippocampal theta oscillations to running and a forelimb-dominated sensorimotor behavior"

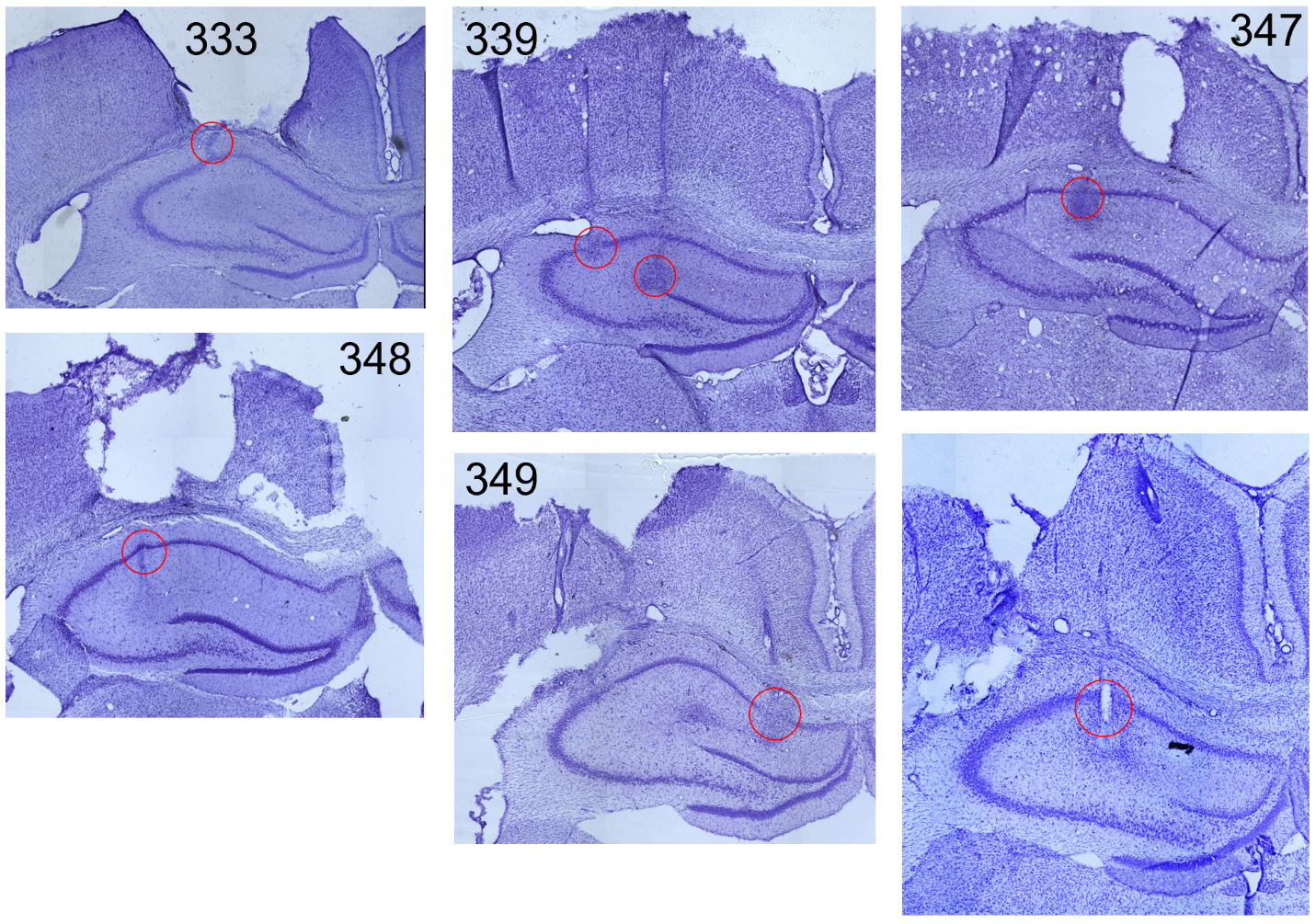

Supplementary Figure 1. Histology (Nissl Stain) centered on the dorsal hippocampus indicating electrode tracks and estimated electrode location (red circle) for each rat.

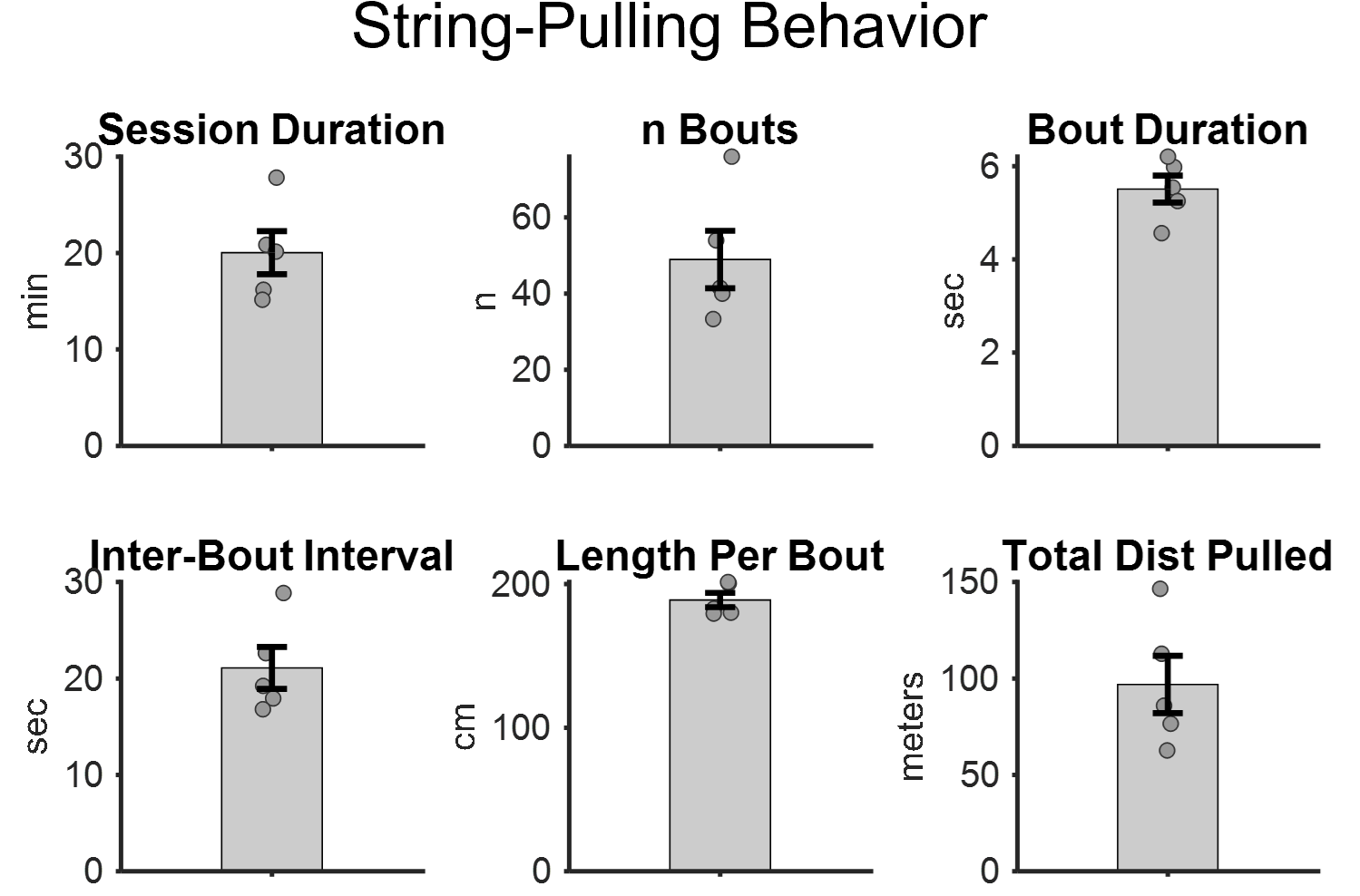

Supplementary Figure 2 String-pulling performance during neural recording sessions. Each subplot indicates a feature of the string-pulling task for each rat (n=5). Session Duration indicates the mean duration of the string-pulling behavior during the recording session. N Bouts indicates the number of string-pulling bouts (where the animal was pulling continuously). Bout Duration is the mean duration of each bout. Inter-bout interval is the mean time between each pulling bout. Length per bout is the length of string pulled per bout. Total Dist Pulled is the mean length of string pulled during any given recording session.
